# Parent of origin gene expression in the bumblebee, *Bombus terrestris*, supports Haig’s kinship theory for the evolution of genomic imprinting

**DOI:** 10.1101/2020.01.17.909168

**Authors:** Hollie Marshall, Jelle S. van Zweden, Anneleen Van Geystelen, Kristof Benaets, Felix Wäckers, Eamonn B. Mallon, Tom Wenseleers

**Author notes:** **Corresponding authors:** Eamonn Mallon.

## Abstract

Genomic imprinting is the differential expression of alleles in diploid individuals, with the expression being dependent upon the sex of the parent from which it was inherited. Haig’s kinship theory hypothesizes that genomic imprinting is due to an evolutionary conflict of interest between alleles from the mother and father. In social insects, it has been suggested that genomic imprinting should be widespread. One recent study identified parent-of-origin expression in honeybees and found evidence supporting the kinship theory. However, little is known about genomic imprinting in insects and multiple theoretical predictions must be tested to avoid single-study confirmation bias. We, therefore, tested for parent-of-origin expression in a primitively eusocial bee. We found equal numbers of maternally and paternally biased expressed alleles. The most highly biased alleles were maternally expressed, offering support for the kinship theory. We also found low conservation of potentially imprinted genes with the honeybee, suggesting rapid evolution of genomic imprinting in Hymenoptera.

**Impact summary:** Genomic imprinting is the differential expression of alleles in diploid individuals, with the expression being dependent upon the sex of the parent from which it was inherited. Genomic imprinting is an evolutionary paradox. Natural selection is expected to favour expression of both alleles in order to protect against recessive mutations that render a gene ineffective. What then is the benefit of silencing one copy of a gene, making the organism functionally haploid at that locus? Several explanations for the evolution of genomic imprinting have been proposed. Haig’s kinship theory is the most developed and best supported.

Haig’s theory is based on the fact that maternally (matrigene) and paternally (patrigene) inherited genes in the same organism can have different interests. For example, in a species with multiple paternity, a patrigene has a lower probability of being present in siblings that are progeny of the same mother than does a matrigene. As a result, a patrigene will be selected to value the survival of the organism it is in more highly, compared to the survival of siblings. This is not the case for a matrigene.

Kinship theory is central to our evolutionary understanding of imprinting effects in human health and plant breeding. Despite this, it still lacks a robust, independent test. Colonies of social bees consist of diploid females (queens and workers) and haploid males created from unfertilised eggs. This along with their social structures allows for novel predictions of Haig’s theory.

In this paper, we find parent of origin allele specific expression in the important pollinator, the buff-tailed bumblebee. We also find, as predicted by Haig’s theory, a balanced number of genes showing matrigenic or patrigenic bias with the most extreme bias been found in matrigenically biased genes.

## Introduction

Genomic imprinting is the differential expression of alleles in diploid individuals, with the expression being dependent upon the sex of the parent from which it was inherited [1]. Multiple evolutionary theories attempt to explain its existence [reviewed in 2]. The most widely accepted explanation is the kinship theory developed by Haig [3]. This theory predicts genomic imprinting arose due to natural selection acting differently on the matrigenes (maternal alleles) and the patrigenes (paternal alleles) of an individual for given processes. For example, in a polyandrous mating system with maternal care (e.g. mammals), patrigenes are predicted to be subject to selection pressures which increase resource allocation from the mother at the expense of siblings. Whereas matrigenes in this scenario are predicted to be selected for equal resource distribution amongst offspring.

The majority of support for this theory comes from studies based on mammals and flowering plant systems [2]. However, it has been suggested haplodiploid social insects can provide an ideal system to independently test Haig’s kinship theory [4]. Colonies of social bees consist of diploid females (queens and workers) and haploid males created from unfertilised eggs. This along with their social structures allows for novel predictions of Haig’s theory.

Research exploring parent-of-origin effects in social insects has focused on the behavioural and physiological outputs of genetic crosses. In the Argentine ant (*Linepithema humile*) paternal effects were observed in care-giving associated behaviours and in sex allocation of offspring [5, 6]. Paternal effects on dominance and stinging behaviour have also been observed in crosses of European and Africanized honeybees [7]. Additionally, Oldroyd *et al*. [8] found a parent-of-origin effect of increased ovary size in honeybees but could not definitively determine which parent this effect was driven by.

More recently, reciprocal crosses and next generation sequencing technologies have been used to identify genes with parent-of-origin allele specific expression patterns in honeybees [9, 10]. Both groups used RNA-Seq to study parent-of-origin gene expression in hybrid crosses of honeybee sub-species. The logic of their test was that as honeybee queens are multiply mated, matrigenes can occur in half sisters and therefore should be selected to moderate worker reproduction in queenless colonies. Patrigenes, on the other hand will not be in half sisters and will be selected to reproduce at any cost [10]. Therefore, the prediction is that parent of origin allele specific expression should exist in honeybees and patrigenic expression will dominate [10]. Surprisingly, Kocher *et al*. [9] found a matrigenic bias in gene expression however, it was later shown that sub-species incompatibility effects influenced the results obtained [11]. Showing support for the kinship theory, Galbraith *et al*. [10] found greater patrigenic expression in reproductive workers compared to sterile workers, with increased patrigenic expression in the reproductive tissues.

The lack of agreement between these studies weakens their support for Haig’s theory. Another weakness of the evidence is that it is limited to only one species and tests only one prediction from the many predictions made [4] for Haig’s theory and genomic imprinting’s role in social insect biology.

To test the robustness of Haig’s kinship theory we present gene expression data (RNA-seq) of reproductive and sterile workers from reciprocal crosses of sub-species of the primitively eusocial bumblebee, *Bombus terrestris*. This species is naturally singly-mated. As such, the predictions for the presence of matrigenic/patrigenic expression bias are different of those predicted for the naturally multiply-mated honeybee [4]. As in Galbraith *et al*. [10] we test the effect of queenless conditions. The same patrigene is present in all sisters (Fig.1), therefore patrigenes should be selected to moderate worker reproduction to reduce the cost to nephews. This is the opposite of the honeybee prediction. Matrigenes, similar to honeybees, have a non-zero probability of being in any given sister (Fig.1), therefore matrigenes should also be selected to moderate worker reproduction, although possibly at a lower level that patrigenes. Therefore we make three predictions; 1) that parent-of-origin allele specific expression exists in bumblebees, 2) that parent-of-origin allele specific expression should be balanced between genes biased patrigenically and matrigenically, with perhaps a slight surplus of matrigenically expressed genes due to the decreased probability of a matrigene occurring in a nephew as opposed to a son and 3) that genes showing parent-of-origin expression will be enriched for reproductive related processes.

**Figure 1:**
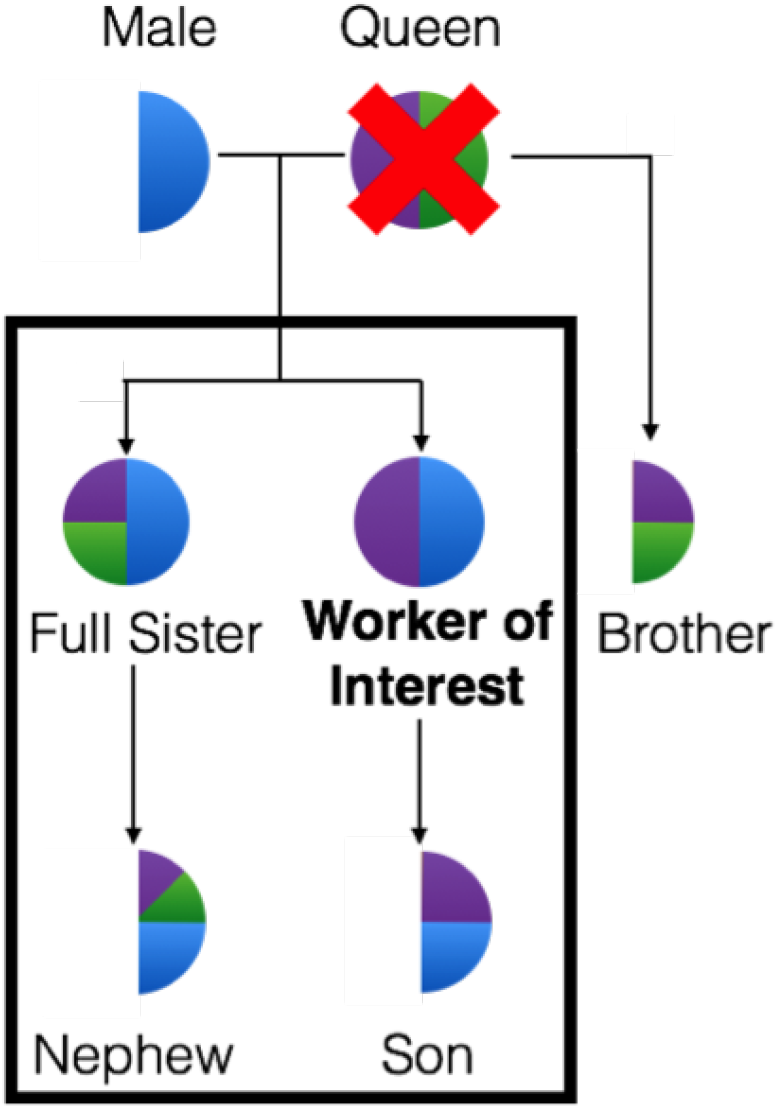
Parental genome inheritance probabilities under queenless conditions in bumblebees. Schematic adapted from Drewell *et al*. [12]. As the bumblebee is singly mated, there is only one patrigene (blue) shared between all workers. This means there is an equal probability of the patrigene ending up in the worker’s son or nephew. The worker of interest has inherited the purple matrigene. Her sister has a 50% change of inheriting the purple or the green matrigene. For the purple matrigene, there is a 50% chance of ending up in the son but a 25% chance of ending up in the nephew.

## Results

### Differential expression between reproductive castes

Ten bases were trimmed from the 3’ end of each worker RNA-seq read due to base bias generated by the Illumina protocol [13]. The mean number of uniquely mapped reads to the maternal genomes was 91.31% ± 0.57% (mean ± standard deviation) for the head samples and 91.53% ± 1.86% for the abdomen samples. This equated to a mean of 13,602,926 ± 2,774,040 and 14,181,003 ± 1,303,486 uniquely mapped reads respectively (supplementary 1.0.1). The mean number of uniquely mapped reads to the paternal genomes was 91.50% ± 0.68% (mean ± standard deviation) for the head samples and 91.70% ± 1.83% for the abdomen samples. This equated to a mean of 13,630,611 ± 2,776,779 and 14,208,273 ± 1,307,054 uniquely mapped reads respectively (supplementary 1.0.1).

Tissue type explains the majority of variation within all of the RNA-seq samples, followed by reproductive status; either reproductive or sterile (supplementary 2.0: Fig.S1). Following differential expression analysis a total of 3,505 genes were up-regulated in the abdomen of reproductive workers compared to sterile workers and 4,069 genes were down-regulated (q<0.01) (supplementary 1.0.2).

The enriched GO terms for the differentially expressed genes between reproductive and sterile castes in the abdomen included mostly regulatory processes but also "*reproduction*" (GO:0000003) (supplementary 1.0.3). Enriched GO terms associated specifically with up-regulated genes in reproductive workers in the abdomen also included: "*reproduction*" (GO:0000003) and "*DNA methylation*" (GO:0006306) (supplementary 1.0.4), these terms were not found in the enriched GO terms for genes up-regulated in sterile workers (supplementary 1.0.5).

Considerably fewer genes were differentially expressed in the head samples; 86 up-regulated genes in reproductive compared to sterile workers and 41 down-regulated genes (q<0.01) (supplementary 1.0.6). The majority of the GO terms associated with these differentially expressed genes involved biosynthetic processes (supplementary 1.0.7). Up-regulated genes in the head tissue of reproductive workers also included "*reproduction*" (GO:0000003) whereas the up-regulated genes in the head tissue of sterile workers consisted of mostly metabolic processes (supplementary 1.0.8 and 1.0.9).

### Parent-of-origin gene expression

A total of 10,211 genes had a minimum of two SNPs with at least a coverage of five reads each. Of those, 7,508 genes occurred in every cross, worker-type and tissue type. 700 genes had significant maternal/paternal expression bias in both cross-directions for reproductive workers (q <0.05), and 747 were significant for sterile workers. The expression bias was averaged across; tissue type, worker type, family and direction of cross to obtain an extremely conservative expression proportion. The significant genes were then filtered to also have an average maternal expression proportion of >0.6 or <0.4 to give a final confident list of genes showing parent-of-origin expression.

Reproductive workers have 163 genes showing significant parent-of-origin expression (Fig.2) (supplementary 1.1.0). Sterile workers have 170 genes showing significant parent-of-origin expression (Fig.2) (supplementary 1.1.1). There is no significant difference between the number of genes showing maternal expression bias compared to paternal expression bias based on reproductive status (chi-squared test of independence, χ-squared = 0, df = 1, p-value = 1). There is also no difference in the number of genes showing paternal expression bias compared to maternal expression bias in reproductive and sterile workers, assessed independently (chi-squared goodness of fit, reproductive: χ-squared = 1.3804, df = 1, p-value = 0.24, sterile: χ-squared = 1.5059, df = 1, p-value = 0.2198). The most extreme expression bias is seen in the maternally expressed alleles in both castes, with 17 genes showing a maternal expression proportion of >0.9 in both reproductive and sterile workers (Fig.2). There were no genes showing >0.9 paternal expression bias. Additionally we did not find any genes with significant sub-species expression bias.

**Figure 2:**
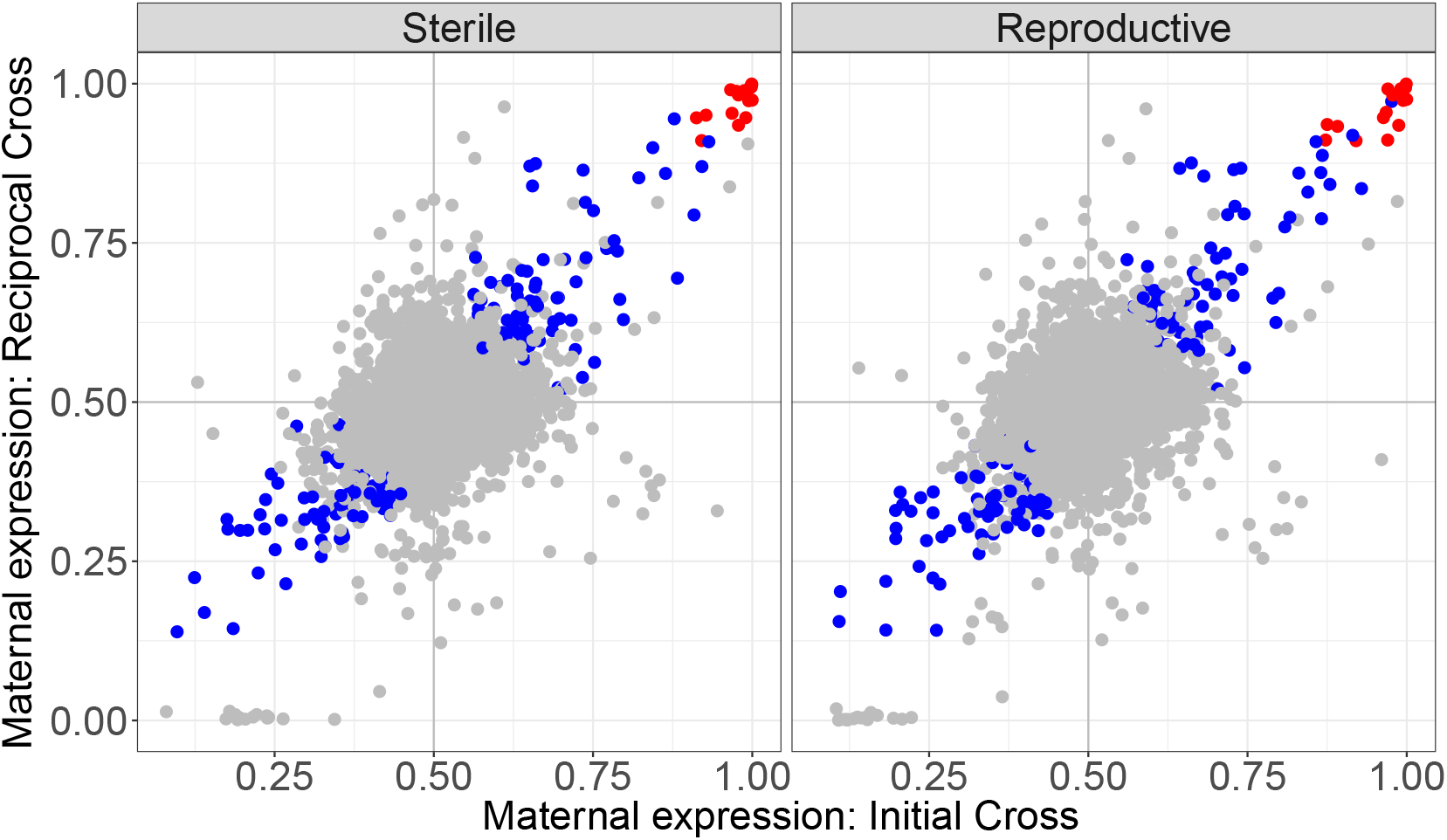
Maternal expression proportion of all genes by worker caste. Each point represents a gene. Blue points are genes with significant parental expression bias (q<0.05 and expression proportion >0.6). Red points are genes with significant parental expression bias (q<0.05) with the proportion of expression >0.9. The top left quadrant of each plot represents genes with a *B. terrestris audax* expression bis, the bottom right quadrant represents gene with a *B. terrestris dalmatinus* expression bias. The top right represents genes with a maternal expression bias and the bottom left represents genes with a paternal expression bias.

Reproductive and sterile workers share a significant number of genes showing parent-of-origin expression with the same parental bias (Fig.3, maternal expression bias: hypergeometric test, p = 9.20 × 10 ^−108^, paternal expression bias: hypergeometric test, p = 7.66 × 10 ^−90^). There were no genes with maternal/paternal bias in one caste which also had the opposite bias in the other caste. Additionally the majority of genes identified as showing parental expression bias show the same bias in both abdomen and head tissue as well as across behaviourally defined castes (dominant and subordinate reproductives and sterile foragers and nurses), see supplementary 2.0: Fig.S2-S7.

**Figure 3:**
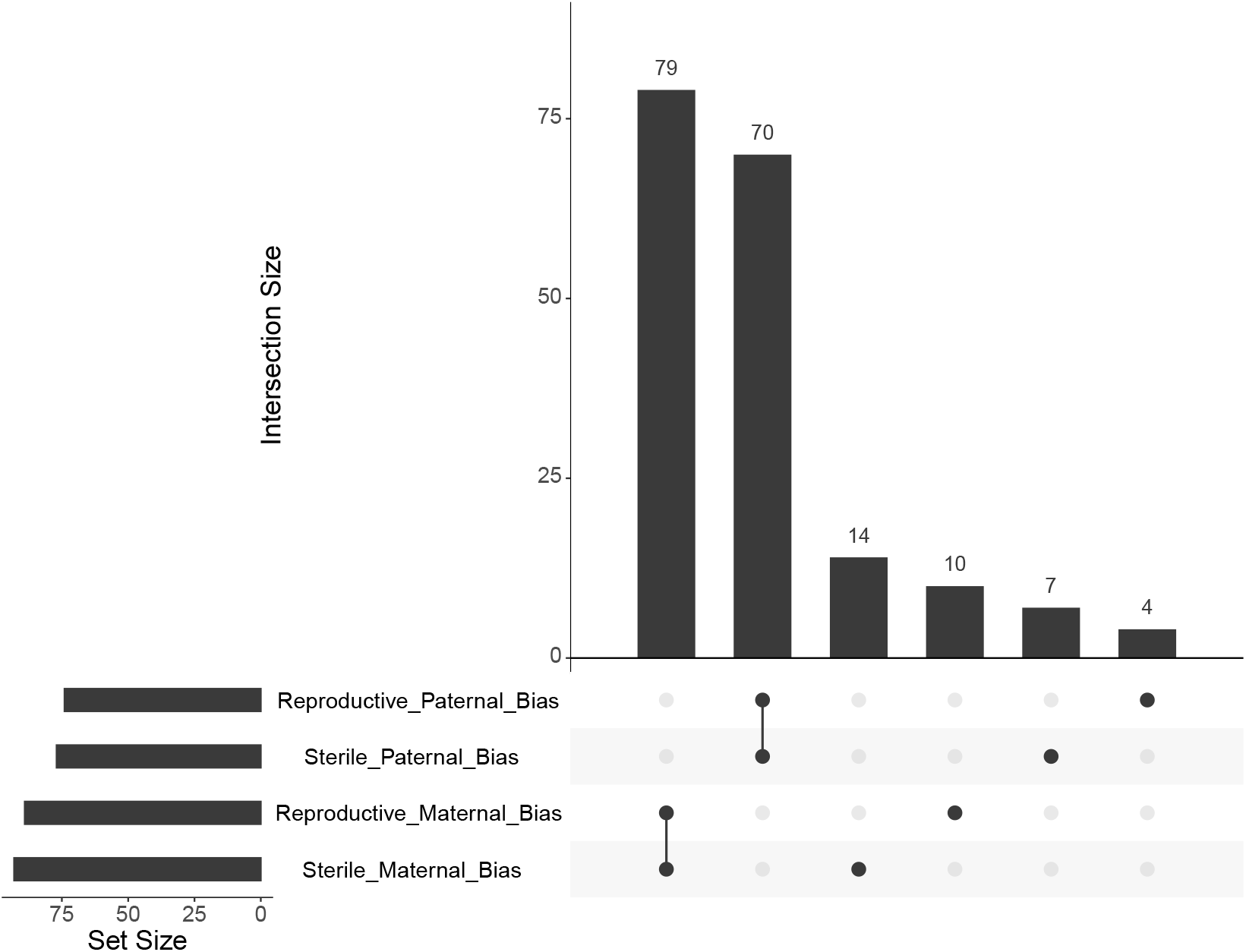
Overlapping genes showing parent-of-origin expression in reproductive and sterile workers. The set size indicates the number of genes in each list. The intersection size shows how many genes the corresponding lists have in common. A single dot refers to the number of genes unique to each list.

Overall genes showing parent-of-origin expression have enriched GO terms for multiple biological processes (supplementary 1.1.2), specifically the GO terms "*negative regulation of reproductive processes*" (GO:2000242) and "*female germ-line sex determination*" (GO:0019099) are enriched. Genes with maternal and paternal bias in both reproductive and sterile workers also have enriched GO terms for multiple biological processes (supplementary 1.1.3 and 1.1.4). Specifically, paternally expressed genes in both reproductive and sterile workers are enriched for the GO term; "*behaviour*" (GO:0007610).

GO terms for genes showing parent-of-origin expression in only reproductive or sterile workers were also enriched for various biological processes (supplementary 1.1.5). Including "*histone ubiquitination*" (GO:0016574) and "*histone H2A monoubiquitination*" (GO:0035518) in reproductive paternally expressed genes.

Genes showing maternal parent-of-origin expression are enriched for genes which are also differentially expressed in head tissue (Supplementary 2.0: Fig.S8, hypergeometric test, p = 0.004). Specifically *Serine Protease Inhibitor 3/4* (LOC100652301) shows maternal expression bias in both reproductive and sterile workers and is up-regulated in the head tissue of reproductive workers.

Genes with paternal parent-of-origin expression do not significantly overlap with differentially expressed genes (supplementary 2.0: Fig.S8, hypergeometric test, p = 0.148). There is also no significant overlap between genes showing parent-of-origin expression bias and differential expression in the abdomen (supplementary 2.0: Fig.S9, hypergeometric test, p = 0.996).

### Honeybee homology

A custom database of putative orthologs was made between *A. mellifera* and *B. terrestris*, as in [14], containing 6,539 genes. 68% of differentially expressed abdominal genes were identified within the database, 43% of differentially expressed head genes and 46% of genes showing parent-of-origin expression bias (supplementary 1.1.6). Gene lists were obtained from Galbraith *et al*. [10], the ortholog database contained 38% of the differentially expressed genes identified between honeybee reproductive worker castes and 53% of the genes found to show parent-of-origin expression bias (supplementary 1.1.6).

There was no significant overlap between the genes identified as showing parent-of-origin expression between both studies (hypergeometric p = 0.64), with only two genes overlapping (supplementary 2.0 Fig.S10). One of these is uncharacterised in both species (honeybee id: LOC552195, bumblebee id: LOC100648162) and the second is a serine protease inhibitor (honeybee id: LOC411889, bumblebee id: LOC100644680). The serine protease inhibitor shows paternal expression bias in honeybees and maternal bias in both bumblebee reproductive and sterile castes. It is not differentially expressed in the honeybee but it shows up-regulation in the abdomen tissue of reproductive bumblebee workers compared to sterile workers. Additionally, [10] identified numerous genes of interest which are involved in reproductive behaviour (*vitellogenin, yolkless, ecdysone-receptor and ecdysone-induced protein*) which show paternal expression bias in honeybees, none of which show significant parent-of-origin expression in the bumblebee (supplementary 2.0 Fig.S11).

There was also no significant overlap between differentially expressed genes identified in head tissue and abdomen tissue between *B. terrestris* reproductive castes with those identified as differentially expressed between reproductive castes of *A. mellifera* from [15] (supplementary 2.0 Fig.S12, head: hypergeometric p = 1, abdomen hypergeometric p = 1).

## Discussion

Using parental genome sequencing and offspring RNA-seq we have identified genes showing parent-of-origin allele specific expression in a primitively eusocial bumblebee species. There was no difference in the number of genes showing maternal or paternal expression bias in either reproductive or sterile workers. The genes showing the highest proportion of expression bias were all maternally expressed in both worker castes. Additionally, reproductive related GO terms were enriched in both maternally and paternally biased genes.

Reproductive and sterile workers were chosen in order to provide a robust test for the kinship theory. Imprinted genes should maintain their expression bias regardless of the current reproductive state of the individual, i.e. higher matrigenic expression compared to patrigenic expression should be present in queenless workers regardless of whether they have become reproductive or remained sterile. Additionally the use of sterile and reproductive workers allowed us to assess differential expression between castes. This meant we could investigate if there was a relationship between differentially expressed genes between reproductive castes and genes showing parent-of-origin expression. A high overlap of differentially expressed genes between reproductive castes with genes showing parent-of-origin expression would suggest a role for imprinted genes in reproductive caste determination.

As in Galbraith *et al*. [10] we also found no significant overlap of paternally expressed genes with differentially expressed genes between reproductive and sterile workers. However, we did find a significant overlap with maternally expressed genes and genes differentially expressed in head tissue between reproductive and sterile castes. This significant overlap should be interpreted cautiously however as only seven of the 103 unique maternally biased genes were differentially expressed in head tissue between reproductive castes. The lack of overlap of differentially expressed genes with paternally expressed genes and the small overlap with maternally expressed genes suggests parent-of-origin expression may not directly influence reproductive status in bumblebee workers.

One of the overlapping genes found to be differentially expressed in head tissue between reproductive castes which also shows maternal expression bias is a serine protease inhibitor. One of the two homologous genes identified as showing parent-of-origin expression in this study and in honeybees was also a serine protease inhibitor. Serine proteases (also know as serpins) have been shown to be involved in insect immunity in various species including; the silkworm *Bombyx mori* [16], another species of silk producing moth *Antheraea pernyi* [17] and the mosquito *Anopheles gambiae* [18]. Most recently a kazal-type serine protease inhibitor has been directly linked to oocyte development in the desert locust *Schistocerca gregaria* [19]. Future work identifying the function of serine protease inhibitors in social insects is needed to better understand the function of parent-of-origin expression of these genes in both *B. terrestris* and *A. mellifera*.

We found genes involved in histone modifications to be paternally expressed in reproductive workers. Histone modifications have been identified as an imprinting mark in plants [20] and thought to be involved in imprinting maintenance in mammals [21]. Histone modifications can alter gene expression by affecting gene accessibility via chromatin [22]. Chromatin modifications have been associated with parent-of-origin expression in the fruit fly *Drosophilia melanogaster* [23]. Galbraith *et al*. [10] also found genes involved in histone modifications showing parent-of-origin expression in the honeybee.

Only two genes were found in common between those found in Galbraith *et al*. [10] as showing parent-of-origin gene expression in the honeybee *A. mellifera* and those identified here in *B. terrestris*. Galbraith *et al*. [10] used ovaries and fat bodies as their tissues samples whereas we selected to test whole head and whole abdomen samples for strong signals of expression bias. Some imprinted genes in mammals are known to be tissue specific, such as *GBR10* which has been found to be maternally expressed in brain and muscle tissue but not in growth plate cartilage [24]. Tissue specificity could account for the lack of concordance in parentally expressed genes found between *B. terrestris* and *A. mellifera*. Additionally 51% of bumblebee genes and 52% of honeybee genes were not present in the homology database created.

Imprinted genes in mammals show much more consistency across species, with mice and humans reportedly sharing around 50 imprinted genes out of around 150 and 100 characterised in each respectively [25]. Additionally, domestic cattle and pigs have been shown to share 14 imprinted genes out of 26 and 18 respectively [26]. However, imprinted genes in plants generally show less conservation, with one study reporting 14% of maternally expressed genes and 29% of paternally expressed genes in *Capsella rubella* show the same imprinting status in *Arabidopsis thaliana*, even though both species belong to the Brassicaceae family [27]. Hatorangan *et al*. [27] suggest the lack of consistency between species could be the result of a historical shift in mating-systems. Given the differences in mating-systems between honeybees and bumblebees, and that variable predictions from the kinship theory apply to each species, rapid evolution of imprinted genes in Hymenoptera is a feasible explanation for the lack of consistency in potentially imprinted genes identified here and in honeybees.

The GO terms associated with both maternally and paternally expressed genes are diverse. It has been suggested imprinted genes can function as a mechanism for plasticity, allowing gene regulation to change depending on environmental conditions by activating the silenced allele and increasing dosage of that gene [28]. Social insects display, sometimes extreme, phenotypic plasticity, where multiple discrete phenotypes (castes) can arise from a single genome within a colony. In some species this is genetically determined [29], however there is growing evidence epigenetic factors may play a role in caste determination in some species [14, 30, 31]. Matsuura *et al*. [32] modelled a genomic imprinting mediated caste determination system in the termite *Reticulitermes speratus* and found this better explained the influence of parental phenotype on offspring than a purely genetic model. Given the diversity of genes found here showing both maternal and paternal expression bias we believe, along with Matsuura [33], that further experimental investigation into the role of genomic imprinting in caste determination in social insects is needed.

The identification of genes showing parent-of-origin expression in this study lays the ground work for future research to identify potential epigenetic mechanisms of allele specific expression in social insects. Genes showing allele specific expression and DNA methylation have been previously identified in *B. terrestris* [34], and genes involved in the reproductive process have been shown to be differentially methylated between reproductive castes [14, 35]. DNA methylation is the mechanism by which some genes are imprinted in mammals and plants [36] and so investigation of parent-of-origin methylation in *B. terrestris* may be fruitful.

These results provide support for Haig’s kinship theory. We found as predicted a balanced number of genes showing matrigenic or patrigenic bias. Also as predicted, the most extreme bias was found in matrigenically biased genes. This is novel, independent support for this important evolutionary theory. The results of this study create a base for many future avenues of research including; gene function analysis of serine protease inhibitors in Hymenoptera, epigenetic mechanisms of imprinting in insects and imprinted genes as a mechanism for plasticity, caste determination and social evolution.

## Methods

### Sample collection

Reciprocal crosses of *B. terrestris dalmatinus* (native to southern Europe) and *B. terrestris audax* (native to the UK) were carried out by Biobest, Leuven. Reciprocal crosses allow subspecies-of-origin effects (i.e. the effect of genotype) to be disentangled from parent-of-origin effects [9, 10]. In order to obtain enough successful colonies multiple males and females for each cross (Fig. 4) were released into cages to mate. Once mating had occurred the male and female were removed. The male was immediately frozen at −80^◦^C and the female was placed in cold conditions for eight weeks to induce diapause. Ten matings were carried out for each cross.

**Figure 4:**
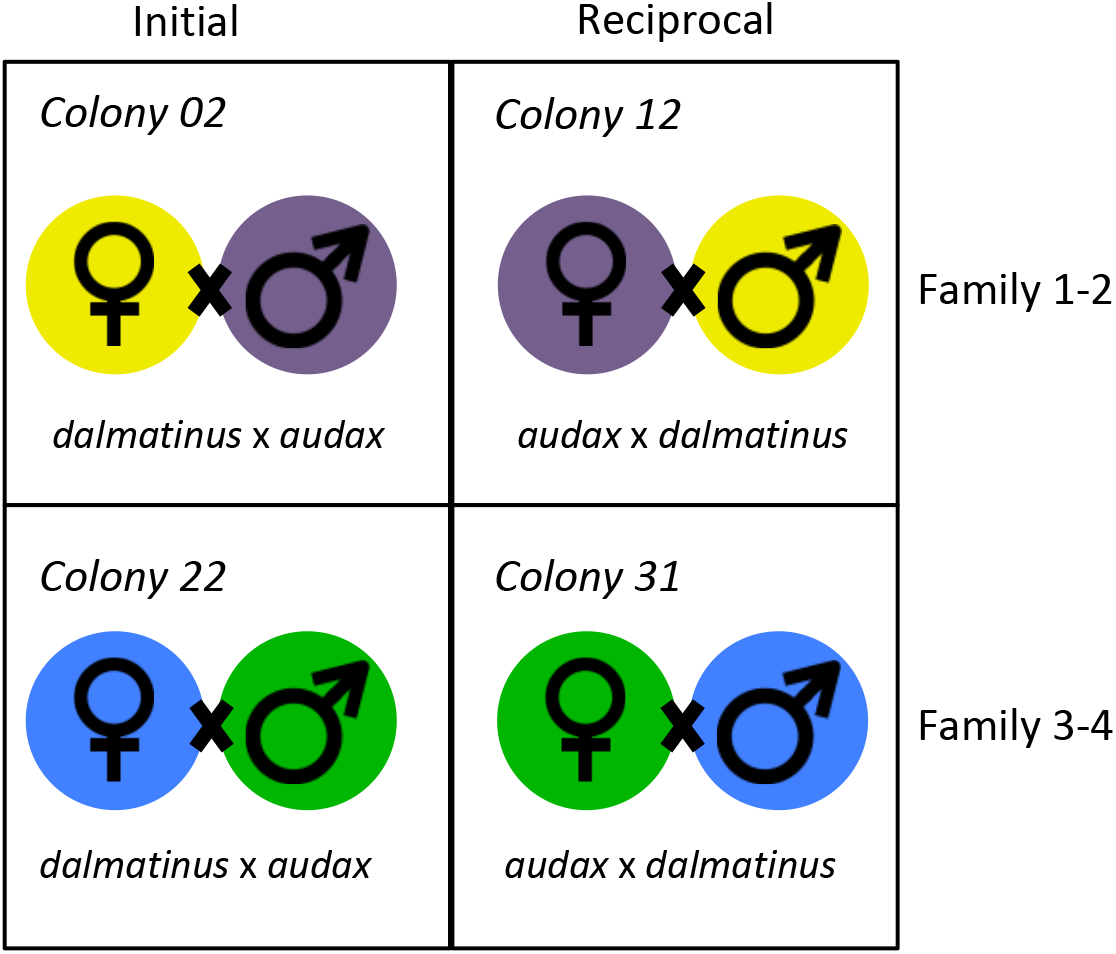
Graphic display of the family-wise reciprocal crosses carried out. Each colour refers to related individuals, i.e. the queen from colony 02 is the sister of the male used in colony 12. This design reduces genetic variability between the initial and reciprocal crosses.

Four successful colonies (one of each cross direction) from two ‘families’ (Fig.4) were housed at the University of Leuven and kept in 21°C with red light conditions, they were fed *ad libitum* with pollen and a sugar syrup. Callow workers were labelled with numbered disks in order to determine age and allow behaviour to be recorded. Once each colony contained approximately 30 workers the queen was removed. The colonies were then filmed under queenless conditions for 30mins per day for 14 days in order to score individual behaviour. The following behaviours were used to classify workers: incubating, feeding larvae, inspecting brood cells, building egg cups, ventilation, biting, pushing, egg-laying, egg-eating, foraging, feeding and grooming. Workers were classified based on the frequency of each of the above behaviours as either: sterile foragers, sterile nurses, dominant reproductives or subordinate reproductives (supplementary 1.0.0).

Worker reproductive status was confirmed by ovary dissection, ovaries were scored on a 0-4 scale as in Duchateau and Velthuis [37], entire bodies were then stored at −80°C along with the original queen mothers and male fathers. Workers were selected for sequencing based on their behavioural classification and ovary status. Two of each behavioural type per colony were selected (i.e. two dominant reproductives, two subordinate reproductives, two nurses and two foragers), with the exception of colony 22 (Fig.4) which contained three subordinate reproductives and one dominant reproductive. All sterile and reproductive samples were age matched. This gave a total of 32 samples, 8 per colony, four of each reproductive status, reproductive or sterile. See supplementary 1.0.0 for behavioural and ovary scoring per sample.

### DNA and RNA extraction and sequencing

DNA was extracted from the mother and father of each colony using the Qiagen DNeasy^®^ Blood & Tissue Kit. Thirty-two workers were selected for RNA sequencing. The head and abdomen were dissected and RNA extracted separately for each, using the Qiagen RNeasy^®^ Lipid Tissue Kit, giving 64 total RNA samples. The quality of the DNA and RNA extraction were measured by Nanodrop and Qubit® fluorometer. Whole genome parental DNA was sequenced using 91bp paired-end reads and worker RNA was sequenced using 90bp paired-end reads on an Illumina HiSeq 2000 by BGI, China. Lane effects were minimised for the RNA samples by spreading colony and tissue samples across five lanes.

### Generation of alternative reference genomes

Whole genome sequencing data was checked using Fastqc (v.0.11.05) (www.bioinformatics.babraham.ac.uk/projects/fastqc/) and adapters and low quality reads were trimmed using Cutadapt v.1.11 [38]. Reads were aligned to the bumblebee reference genome (Bter_1.0, Refseq accession no. GCF_000214255.1 [39]) using BWA-mem v.0.7.15 [40] with standard parameters. SNPs were then called using freebayes v.1.1.0 [41] which can account for the difference in ploidy between males and females, a minimum of five reads per SNP were required. Queen SNPs were then filtered so only the homozygous alternative SNPs remained. The subtractbed command from BEDtools v.2.25.0 [42] was then used to create files containing SNPs unique to either the mother/father of each colony. The individual parental SNP files were then used to create alternate reference genomes for each parent using the ‘fasta alternate reference maker’ command in GATK v.3.6 [43].

### Identification of parent-of-origin expression

RNA-Seq data was quality checked and trimmed as above. STAR v.2.5.2b [44] was used to align worker RNA-seq reads to each of that colony’s specific parental genomes with zero mismatches allowed. This ensures any reads containing a SNP will only be matched to the parent that allele was inherited from. Alignment files were then filtered using BEDtools v.2.25.0 [42] so only alignments which contain an informative SNP (a unique SNP from either the mother or father) were kept. Reads were counted for the maternal/paternal alignments, also using BEDtools v.2.25.0 [42] and SNP positions were annotated with a gene ID taken from the Bter_1.0 annotation file (Refseq accession no. GCF_000214255.1) using a custom R script. SNPs which had zero maternal reads in at least one sample were removed completely from the analysis to avoid possible inflation of paternal counts. This would occur if the queen position was mis-called as homozygous with the missing allele matching that of the male [10].

Genes showing parent-of-origin expression were determined using a logistic regression model in R v.3.4.0 (https://cran.r-project.org). Only genes occurring in both cross directions and in both family combinations, with a minimum of two SNPs per gene were analysed, this left a total of 7508 genes. If any gene showed zero reads for paternal counts this was changed to 1 to avoid complete separation. A quasibionimal distribution was also used to account for overdispersion within the data. Fixed factors included the direction of the cross, family and reproductive status (reproductive or sterile). Correction for multiple testing was carried out using the Benjamini–Hochberg method [45]. Genes were determined as showing parent-of-origin expression if the allelic ratio (maternal/paternal) corrected p-value was <0.05 and the parental expression proportion was >0.6.

### Differential expression

All RNA-seq samples were aligned to the reference genome (Bter_1.0, Refseq accession no. GCF_000214255.1 [39]) using STAR v.2.5.2b [44] with standard parameters. HTseq v.0.8.0 [46] was then used to count the number of reads per gene for each sample. Differential gene expression between reproductive and sterile workers for head and abdomen samples was assessed using the DESeq2 package v.1.16.1 [47] in R. DESeq2 allows the incorporation of a general linear model (GLM) to identify differential expression; family, age, weight, direction of the cross, tissue type and reproductive status were all factors. P-values were corrected for multiple testing using the the Benjamini–Hochberg method [45].

### Gene ontology enrichment

GO enrichment analysis was carried out using the hypergeometric test with Benjamini-Hochberg [45] multiple-testing correction in a custom R script, (q <0.05). This script utilised a previously made GO database for the *B. terrestris* genome, Bter_1.0 [48]. GO terms for differentially expressed genes were tested for enrichment against GO terms associated with all genes identified in either the the RNA-seq data from the abdomen or head. Up-regulated genes in either reproductive or sterile workers were tested for GO term enrichment against all differentially expressed genes from the respective tissue type as a background set.

Genes showing parent-of-origin expression were tested for enrichment against GO terms associated with all genes identified in both abdomen and head RNA-Seq data sets. Genes maternally or paternally biased were checked for GO term enrichment against all genes showing parent-of-origin expression as a background set. REVIGO [49] was used to obtain the GO descriptions from the GO identification numbers.

### Comparative analyses

A hypergeometric test was applied to gene lists from the differential expression analysis and the parent-of-origin expression analysis to identify potential enrichment. *B. terrestis* and *A. mellifera* orthologous genes were determined as in Marshall *et al*. [14] and a custom R script was then used to check for overlap between genes identified as showing parent-of-origin expression here and orthologous *A. mellifera* genes identified in Galbraith *et al*. [10].

## Acknowledgements

We thank Biobest N.V. for providing us with the forty bumblebee colonies of the desired reciprocal crosses. Hollie Marshall had a Ph.D-scholarship from the Central England NERC Training Alliance (NERC, UK). Kristof Benaets had a Ph.D.-scholarship of the Flemish Agency for Innovation by Science and Technology (IWT-Vlaanderen). This study was funded in the context of the KU Leuven BOF Centre of Excellence Financing on ‘Eco- and socio-evolutionary dynamics’ (Project number PF/2010/07) and a grant from the Research Foundation-Flanders (FWO-Vlaanderen, grant G.0463.12). E.B.M. was funded by NERC grant NE/N010019/1. This research used the ALICE2 High Performance Computing Facility at the University of Leicester.

## Author Contributions

T.W. and E.B.M. conceived the study. The reciprocal crosses were carried out by Biobest (Westerlo, Belgium) under supervision of F.W. J.S.Z. and K.B. carried out the behavioural observations and lab work. H.M. and T.W. carried out the analyses with input from A.V.G. H.M. and E.B.M. wrote the initial manuscript. All authors contributed to and reviewed the manuscript.

## Data Accessibility

All RNA-seq and whole genome sequencing data is available under NCBI BioProject number PRJNA329487. Custom scripts are available at the following DOI: https://zenodo.org/record/3235636.

