## Supplementary 2 for "Parent of origin gene expression in the bumblebee, *Bombus terrestris*, supports Haig’s kinship theory for the evolution of genomic imprinting"

### Genome-wide search for parent-of-origin allele specific expression in *Bombus terrestris*: Supplementary 2.

#### 2.0: Supplementary Figures

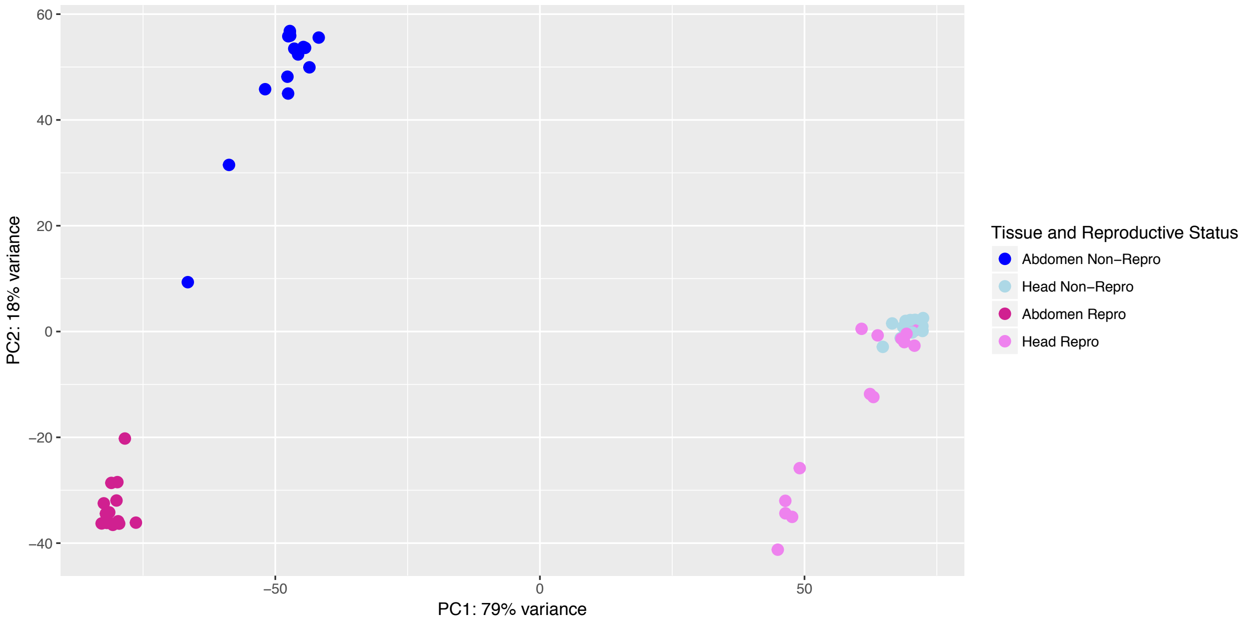


Figure S1: PCA plot based on gene expression data. Samples separate by tissue type along PC1 and by reproductive status along PC2.

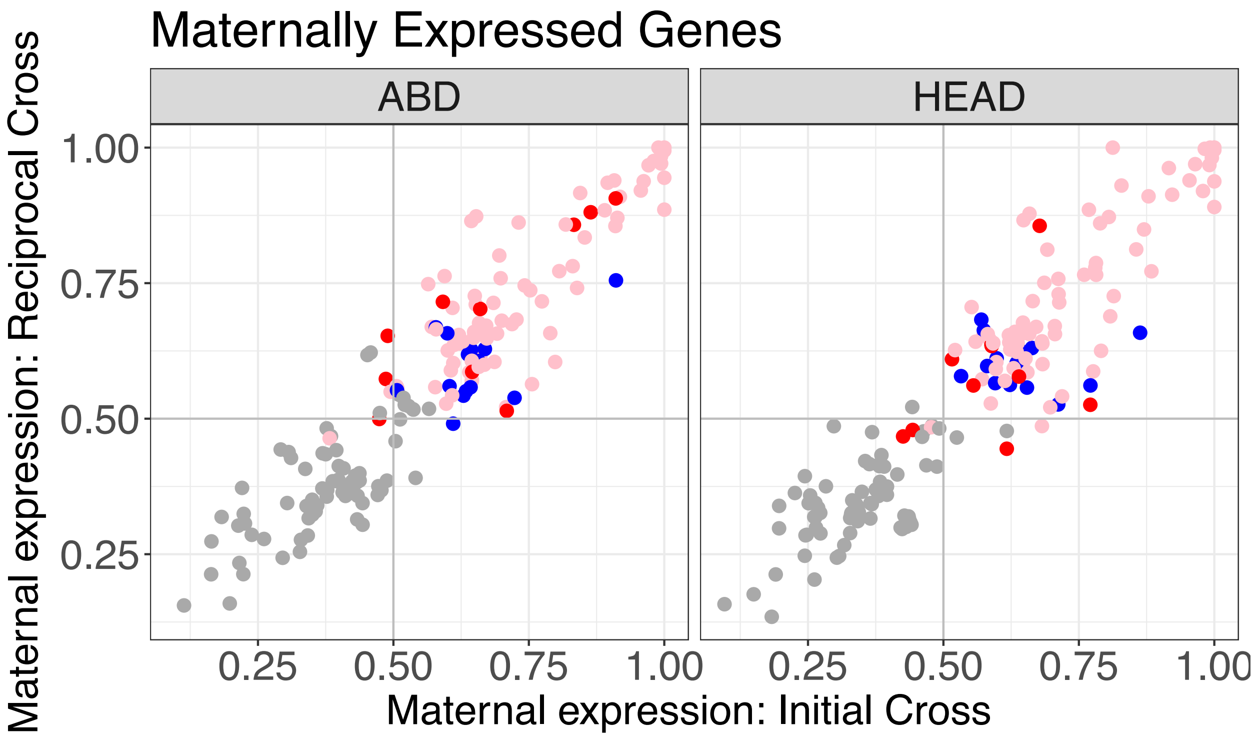


Figure S2: The proportion of maternal expression for each tissue type of all genes found to show significant parental expression bias. ABD refers to abdominal tissue and HEAD refers to head tissue. Each dot represents a gene, the pink dots highlight significantly maternally expressed genes in both sterile and reproductive workers. The red dots are genes showing significant maternal expression bias in reproductive workers only and the blue dots show significant maternal expression bias in sterile workers only.

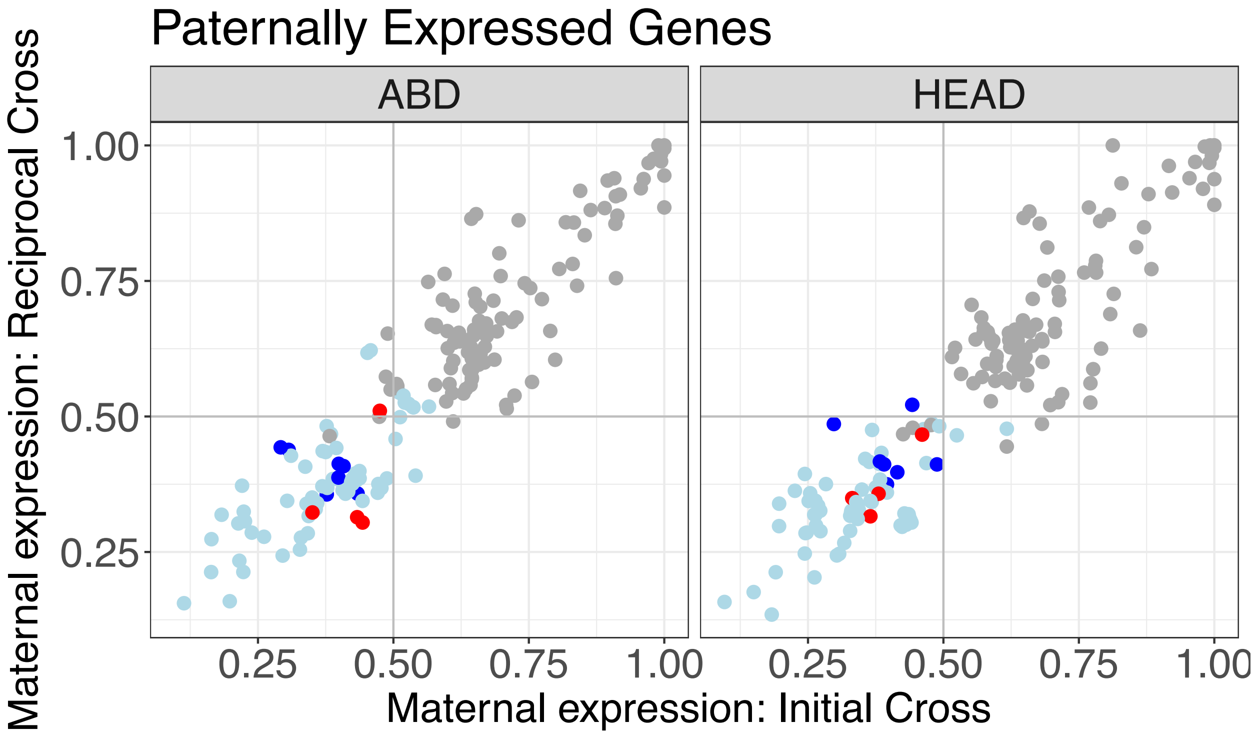


Figure S3: The proportion of paternal expression for each tissue type of all genes found to show significant parental expression bias. ABD refers to abdominal tissue and HEAD refers to head tissue. Each dot represents a gene, the light blue dots highlight significantly paternally expressed genes in both sterile and reproductive workers. The red dots are genes showing significant paternal expression bias in reproductive workers only and the blue dots show significant paternal expression bias in sterile workers only.

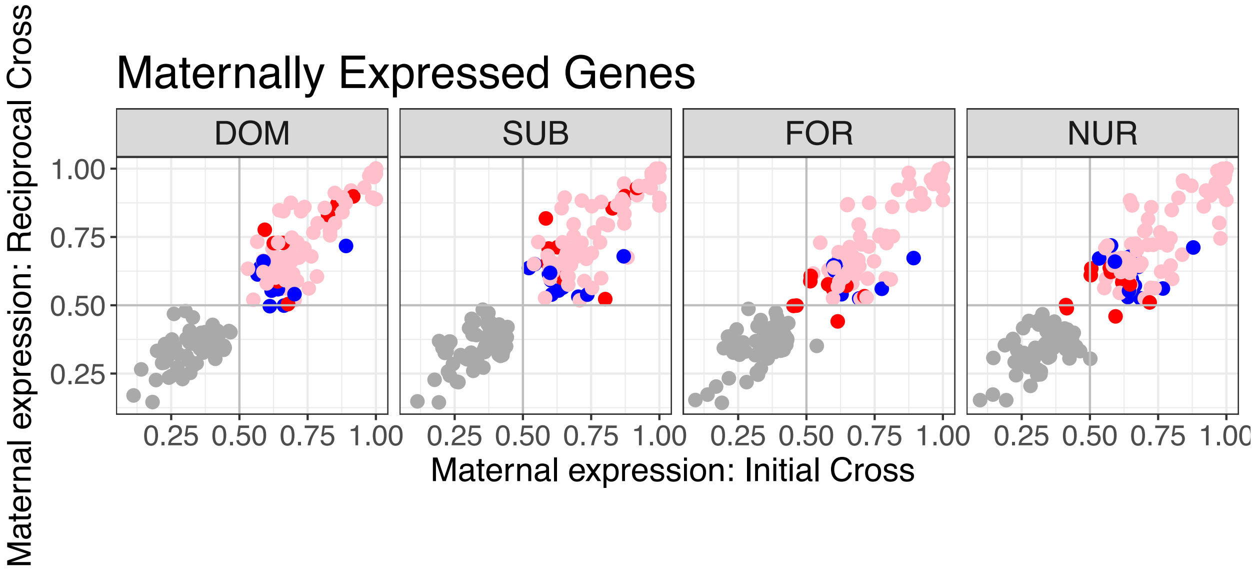


Figure S4: The proportion of maternal expression for each behavioural reproductive type for all genes found to show significant parental expression bias. DOM represents dominant reproductives. SUB represents subordinate reproductives. FOR represents sterile foragers. NUR represents sterile nurses. Each dot represents a gene, the pink dots highlight significantly maternally expressed genes in both sterile and reproductive workers. The red dots are genes showing significant maternal expression bias in reproductive workers only and the blue dots show significant maternal expression bias in sterile workers only.

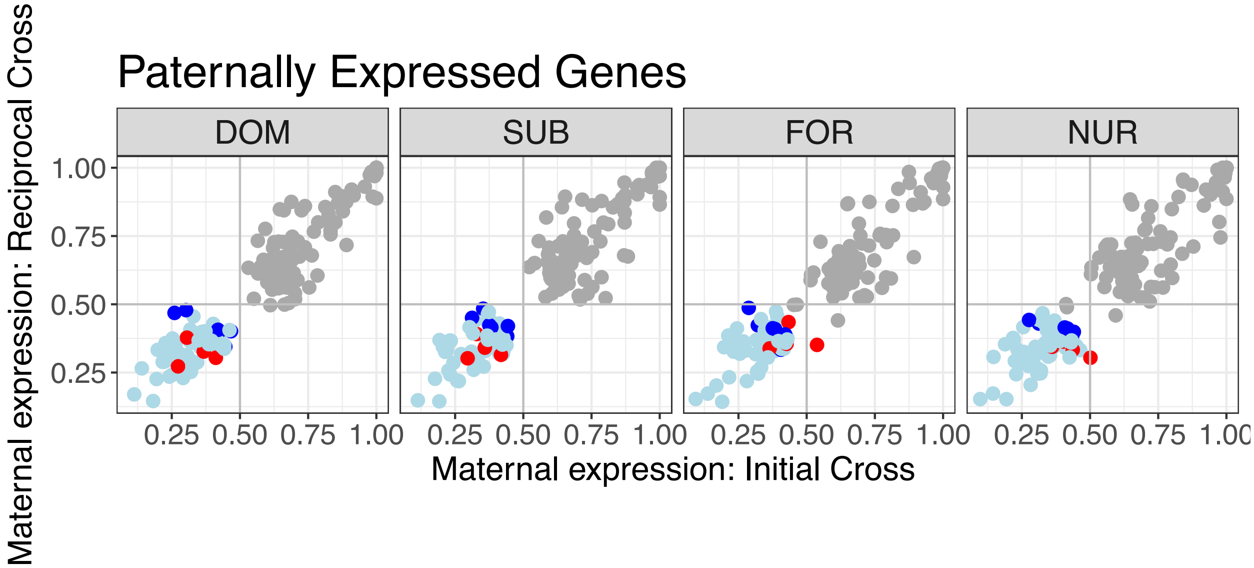


Figure S5: The proportion of paternal expression for each behavioural reproductive type for all genes found to show significant parental expression bias. DOM represents dominant reproductives. SUB represents subordinate reproductives. FOR represents sterile foragers. NUR represents sterile nurses. Each dot represents a gene, the light blue dots highlight significantly paternally expressed genes in both sterile and reproductive workers. The red dots are genes showing significant paternal expression bias in reproductive workers only and the blue dots show significant paternal expression bias in sterile workers only.

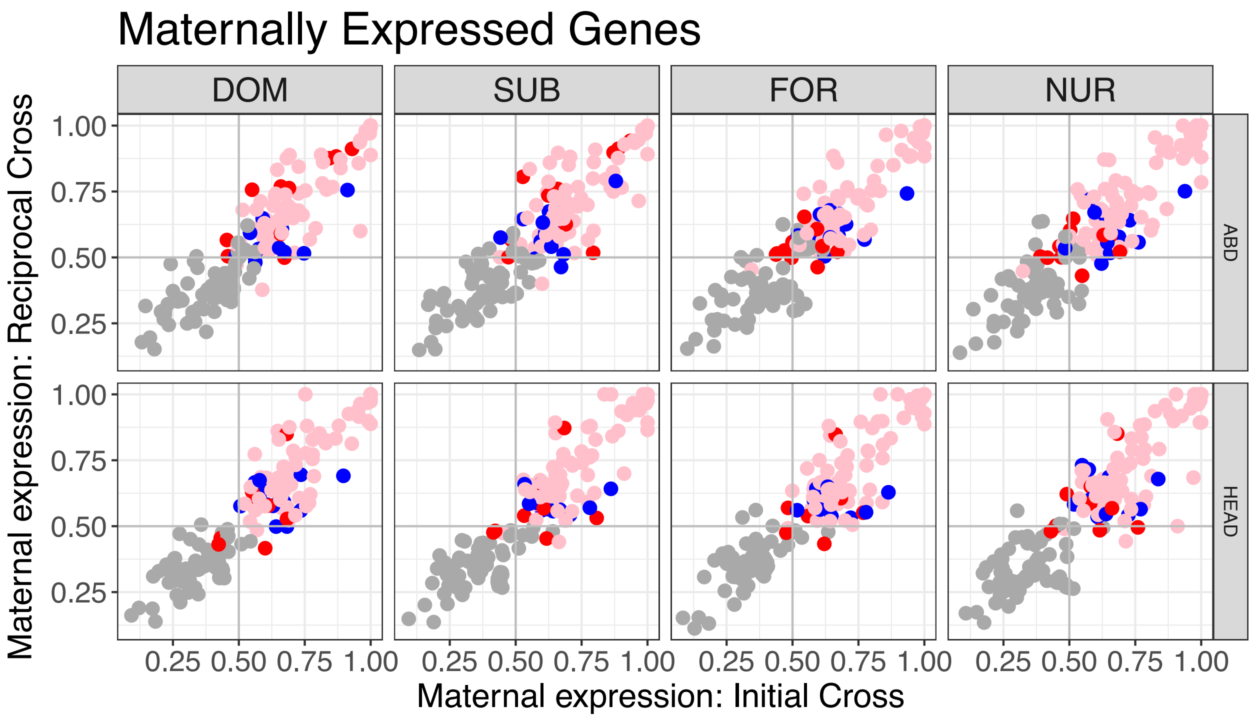


Figure S6: The proportion of maternal expression for each behavioural reproductive type per tissue type for all genes found to show significant parental expression bias. DOM represents dominant reproductives. SUB represents subordinate reproductives. FOR represents sterile foragers. NUR represents sterile nurses. ABD refers to abdominal tissue and HEAD refers to head tissue. Each dot represents a gene, the pink dots highlight significantly maternally expressed genes in both sterile and reproductive workers. The red dots are genes showing significant maternal expression bias in reproductive workers only and the blue dots show significant maternal expression bias in sterile workers only.

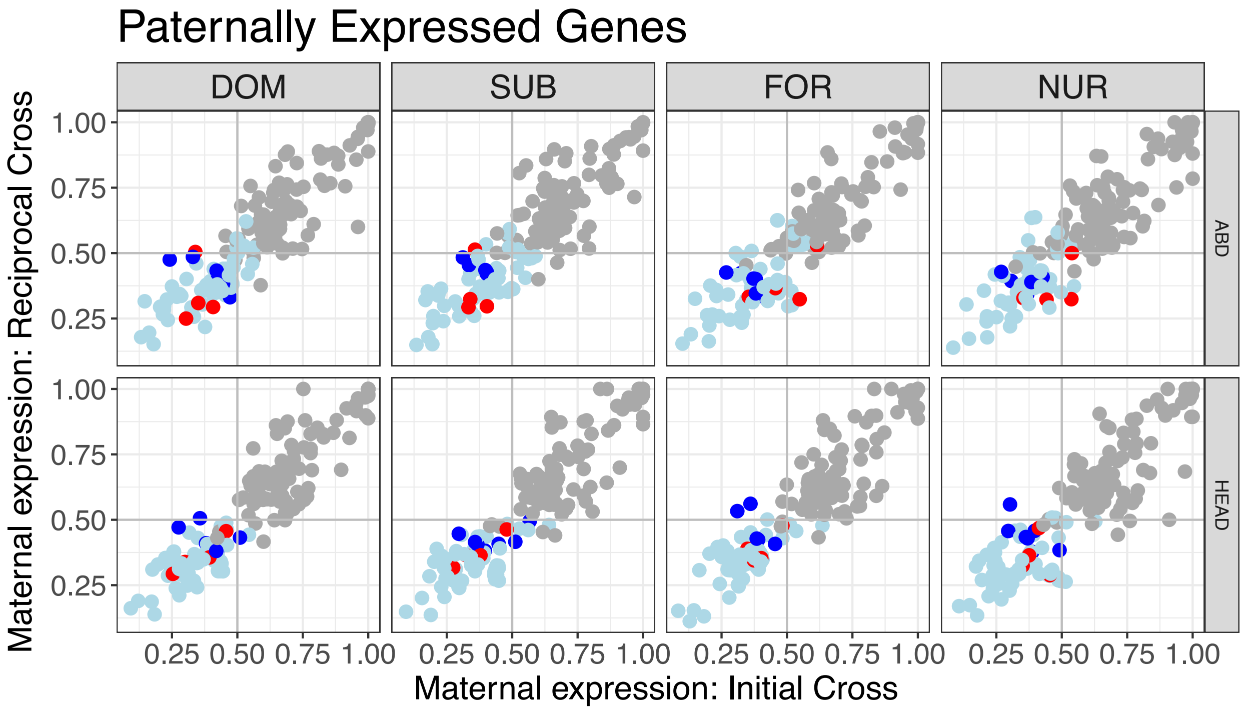


Figure S7: The proportion of paternal expression for each behavioural reproductive type per tissue type for all genes found to show significant parental expression bias. DOM represents dominant reproductives. SUB represents subordinate reproductives. FOR represents sterile foragers. NUR represents sterile nurses. ABD refers to abdominal tissue and HEAD refers to head tissue. Each dot represents a gene, the light blue dots highlight significantly paternally expressed genes in both sterile and reproductive workers. The red dots are genes showing significant paternal expression bias in reproductive workers only and the blue dots show significant paternal expression bias in sterile workers only.

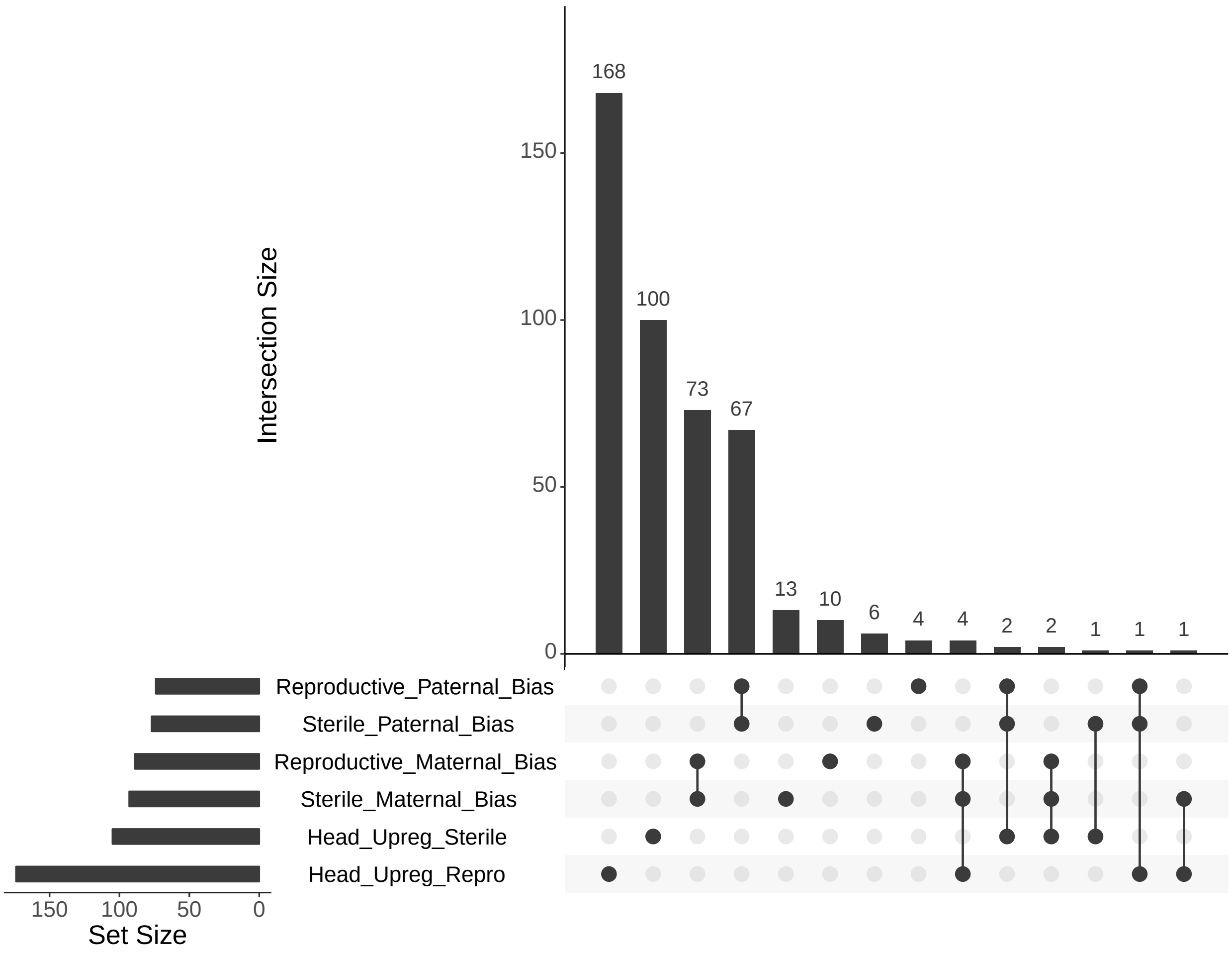


Figure S8: Overlapping genes showing parent-of-origin expression in reproductive (queen-right) and sterile (queen-less) workers with genes showing differential expression in head tissue. The set size indicates the number of genes in each list. The intersection size shows how many genes the corresponding lists have in common. A single dot refers to the number of genes unique to each list. 7/103 genes showing maternal bias are also differentially expressed in the head.

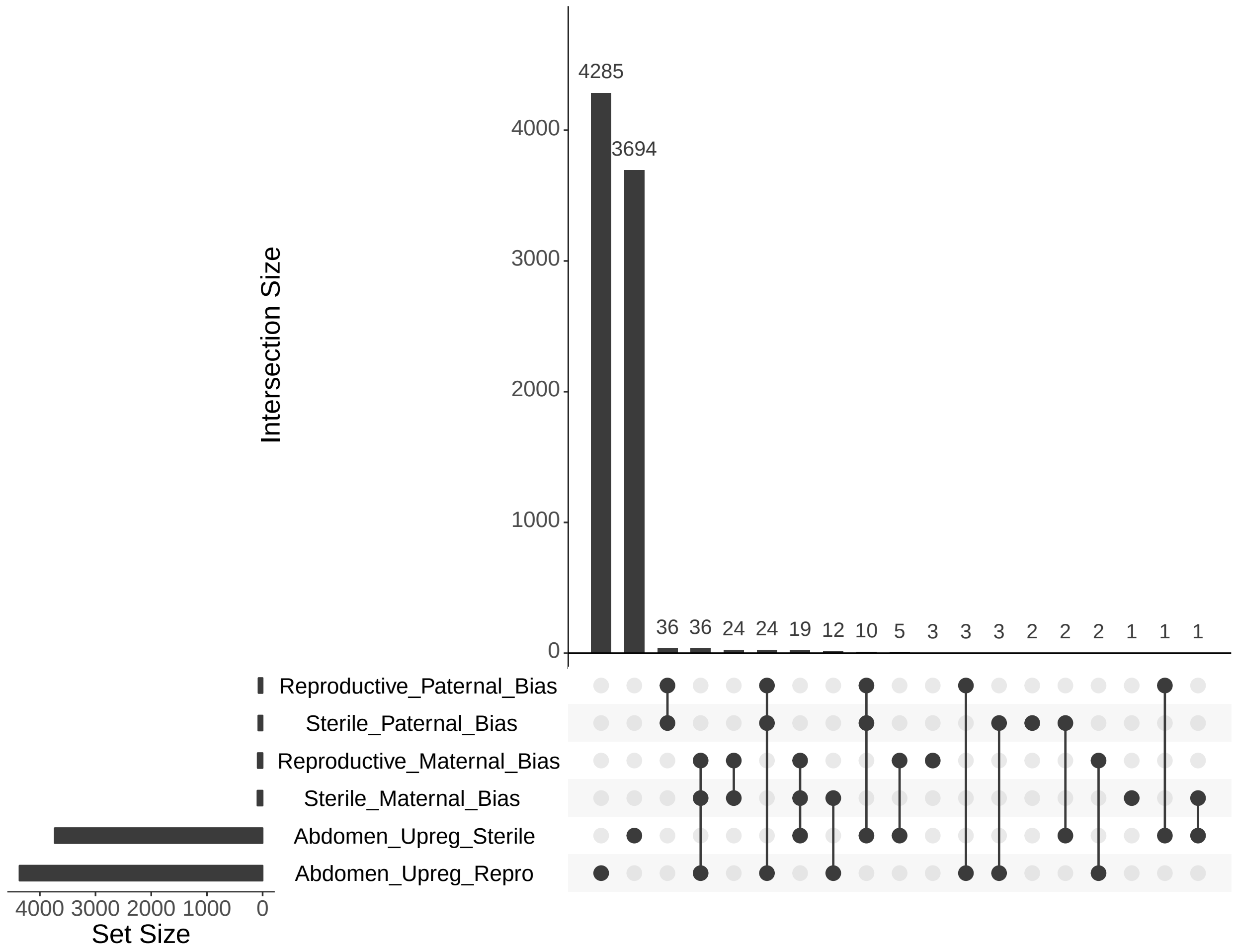


Figure S9: Overlapping genes showing parent-of-origin expression in reproductive (queen-right) and sterile (queen-less) workers with genes showing differential expression in abdomen tissue. The set size indicates the number of genes in each list. The intersection size shows how many genes the corresponding lists have in common. A single dot refers to the number of genes unique to each list.

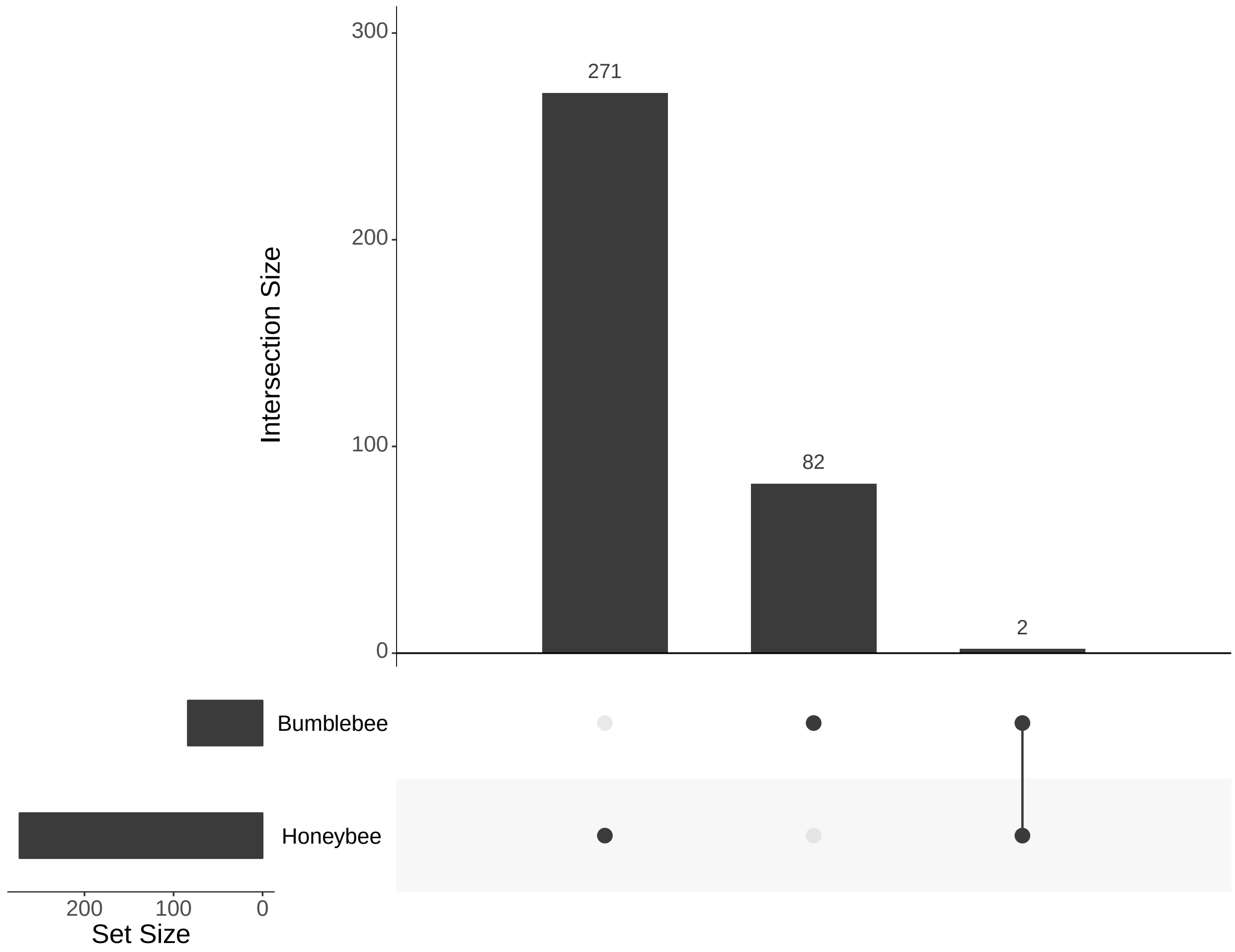


Figure S10: Overlapping genes showing parent-of-origin expression in reproductive (queen-right) and sterile (queen-less) workers of *B. terrestris* and *A. mellifera* (genes identified in Galbraith et al. (2016)). The set size indicates the number of genes in each list. The intersection size shows how many genes the corresponding lists have in common. A single dot refers to the number of genes unique to each list.

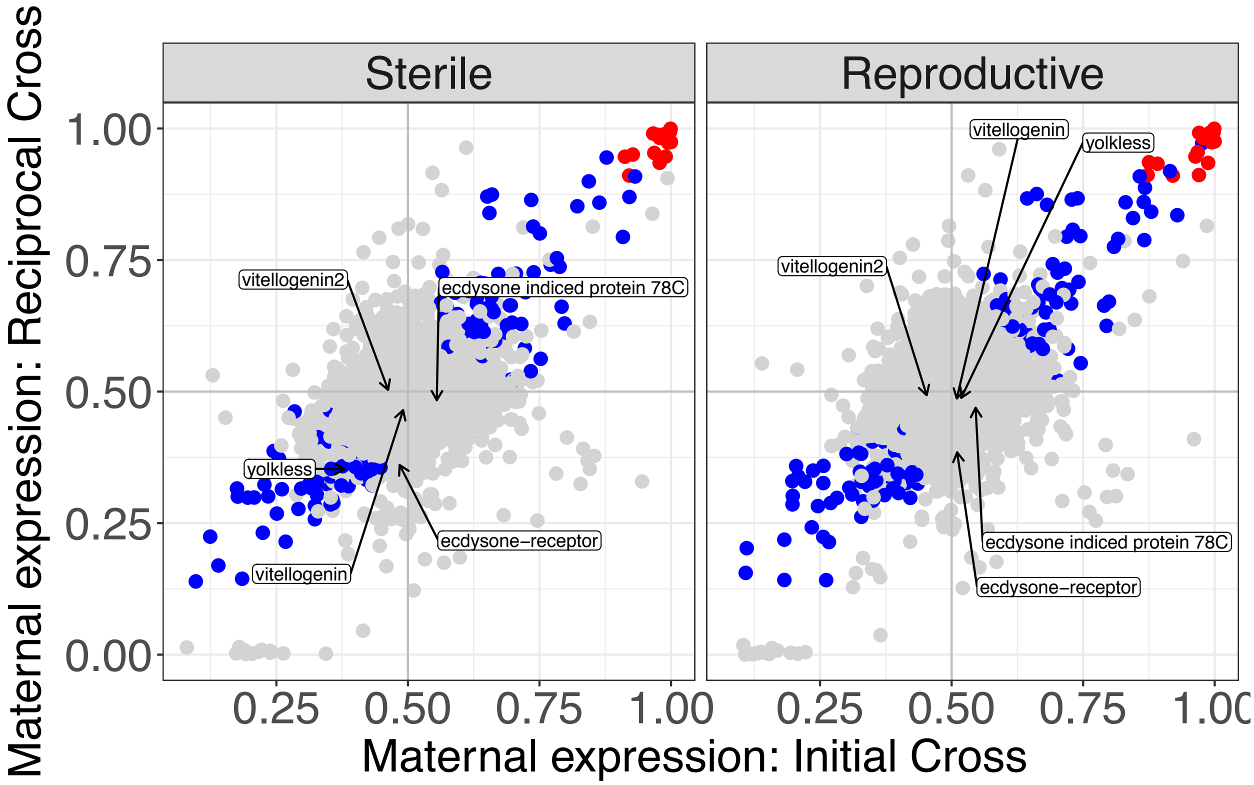


Figure S11: Maternal expression proportion by worker caste, sterile and reproductive. Each point represents a gene. Blue points are genes with significant parental expression bias (q<0.05 and expression proportion >0.6). Red points are genes with significant parental expression bias (q<0.05) with the proportion of expression >0.9. The top left quadrant of each plot represents genes with a *B. terrestris audax* expression bis, the bottom right quadrant represents gene with a *B. terrestris dalmatinus* expression bis. The top right represents genes with a maternal expression bias and the bottom left represents genes with a paternal expression bias. Genes which showed significant paternal expression bias in honeybees (Galbraith et al., 2016) are labelled.

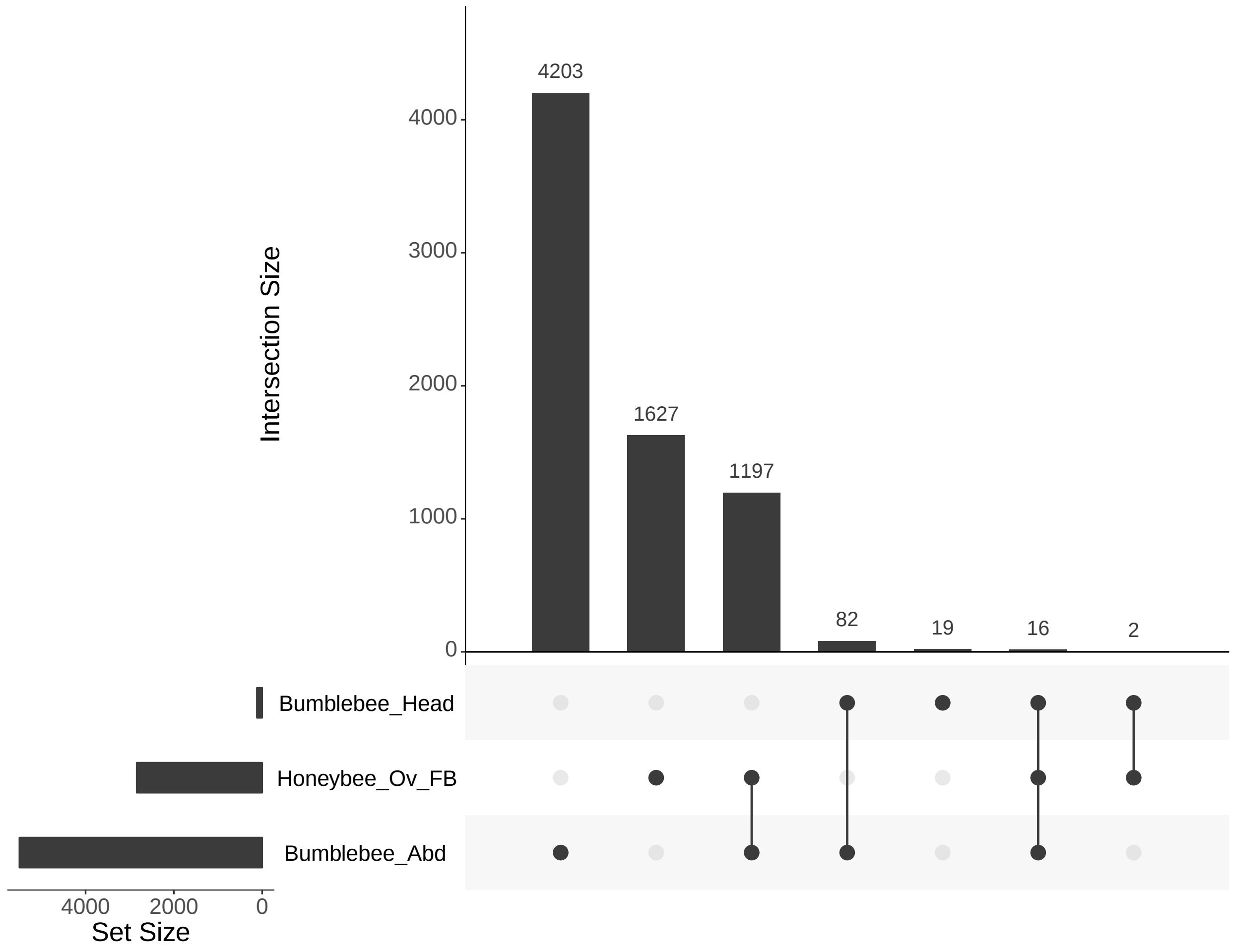


Figure S12: Overlapping genes showing differential expression between reproductive (queen-right) and sterile (queen-less) workers of *B. terrestris* head and abdomen tissue and *A. mellifera* ovary/fatbody tissue (genes identified in Galbraith et al. (2016)). The set size indicates the number of genes in each list. The intersection size shows how many genes the corresponding lists have in common. A single dot refers to the number of genes unique to each list.
